# No title, no theme: The joined neural space between speakers and listeners during production and comprehension of multi-sentence discourse

**DOI:** 10.1101/2019.12.25.884775

**Authors:** Karin Heidlmayr, Kirsten Weber, Atsuko Takashima, Peter Hagoort

## Abstract

Speakers and listeners usually interact in larger discourses than single words or even single sentences. The goal of the present study was to identify the neural bases reflecting how the mental representation of the situation denoted in a multi-sentence discourse (situation model) is constructed and shared between speakers and listeners. An fMRI study using a variant of the ambiguous text paradigm was designed. Speakers produced ambiguous texts in the scanner and listeners subsequently listened to these texts in different states of ambiguity: preceded by a highly informative, intermediately informative or no title at all. Conventional BOLD activation analyses in listeners, as well as inter-subject correlation analyses between the speakers’ and the listeners’ hemodynamic time courses were performed. Critically, only the processing of disambiguated, coherent discourse with an intelligible situation model representation involved (shared) activation in bilateral lateral parietal and medial prefrontal regions. This shared spatiotemporal pattern of brain activation between the speaker and the listener suggests that the process of memory retrieval in medial prefrontal regions and the binding of retrieved information in the lateral parietal cortex constitutes a core mechanism underlying the communication of complex conceptual representations.

## 1. Introduction

Telling a story and listening to it are highly complex neurocognitive activities. To successfully produce or comprehend a story, i.e., a *discourse*, numerous neurocognitive processes are involved. These include processing the articulatory motor programs or the acoustic input of the speech signal, processing the linguistic information at the phonological, semantic and syntactic level, and the conceptual information at the discourse level. But this is not all. Importantly, to understand a story’s meaning, information needs to be integrated over an extended period of time to build a coherent *situation model*, i.e., a mental representation of the situation denoted in a text (Zwaan and Radvansky 1998; Zwaan and Singer 2003). All these processes are required to allow for the speaker and listener to process discourse in its full scope, from the surface structure to the situation model. Moreover, the notion of *parity of representations* between production and comprehension conveys the idea that the same linguistic knowledge and situation model are at stake when an utterance is produced and when it is comprehended (Pickering and Garrod 2004; Pickering and Garrod 2006). Although communication of complex content relies on language use beyond the single word or sentence level, the neural basis underlying the shared conceptualization between interlocutors on the discourse level is not yet well known. Consequently, the present study attempts to shed light on two specific processes, namely the brain activity reflecting the construction of a situation model as well as the brain activity of speakers and listeners related to their engagement in a discourse-level parity of representations.

To understand the meaning of the successive utterances in a discourse, activation in language processing regions of the brain is required. Within the core left-lateralized fronto-temporo-parietal language network, the inferior frontal cortex is a key region for linguistic unification processes, and superior and middle temporal regions play a major role in retrieving language-relevant information from memory (Xiang et al. 2010; Hagoort 2016, 2017). However, in order to construct a coherent situation model of the discourse, more information needs to be retrieved and integrated. Indeed, more extended neural activation is usually found when processing coherent discourse, involving medial frontal as well as medial and lateral parietal regions, among others (Ferstl et al. 2008; Binder 2016; Hagoort 2017; Hasson et al. 2018). Some previous studies and meta-analyses have attempted to identify the functional role of these areas specifically in coherent discourse processing. Importantly, each of these regions is involved in other cognitive tasks as well, and their functions may hence not be limited to the neurocognitive processes listed below. In discourse processing, the left dorsomedial prefrontal cortex (dmPFC) has been argued to play a role in coherence building (Ferstl et al. 2008), inferencing (Kuperberg et al. 2006), and in the top-down retrieval of semantic information stored in temporo-parietal cortices (Binder and Desai 2011). The ventromedial prefrontal cortex (vmPFC) is a crucial region to activate event schemas - distributed neocortical representations from long term memory – and to make them available for the integration of incoming information into a coherent situation model (Nieuwenhuis and Takashima 2011; van Kesteren et al. 2012; Gilboa and Marlatte 2017). The medial parietal cortex, the posterior cingulate cortex (PCC) and precuneus, are involved in integrating information on a large timescale (Hasson et al. 2015; Baldassano et al. 2017), and in episodic memory retrieval and self-centered mental imagery (Cabeza and Nyberg 2000; Cavanna and Trimble 2006). Moreover, in the lateral parietal cortex, the angular gyrus (AG) plays a critical role in semantic unification (Menenti et al. 2008; Hagoort 2017) and in conceptual combination (Price et al. 2015). Finally, activation in the right inferior frontal gyrus (rIFG) has also been associated with the formation of a situation model of an ongoing discourse (Menenti et al. 2008). Accordingly, it can be concluded from previous research that a network of functionally heterogeneous neural regions is involved in processing discourse and constructing a coherent situation model. Here, we were interested in identifying the neural activity related to the construction of the situation model more specifically. Hence, the first goal of the present study was to isolate the brain activity related to situation model construction. To do so, we placed texts with identical wording in different contexts. Each context influenced the ambiguity of the text with respect to the situation described, and hence the ease of constructing a coherent situation model. Moreover, to date, it is an open question to which degree the involved neurocognitive processes are shared between speakers and listeners. Importantly, it has previously been suggested that linguistic alignment between speakers and listeners draws on linguistic processes and semantic memory, whereas the sharing of the situation model draws on episodic memory processes (Pickering and Garrod 2013), which seem to activate different structures of the brain. Thus, the second goal of the present study was to identify the neural bases of the sharing of situation models across speakers and listeners.

To investigate these two questions, we implemented a variant of the ambiguous text paradigm (Dooling and Lachman 1971; Bransford and Johnson 1972). Ambiguous texts consist of a series of grammatically correct but semantically vague sentences. From these sentences, it is difficult to form a meaningful situation model in the absence of an informative context. In contrast, with an informative, disambiguating context (e.g., picture or verbal description), the comprehension and memorization of the conceptual content increases (Dooling and Lachman 1971; Bransford and Johnson 1972; Wiley and Rayner 2000; Wahlberg and Magliano 2004; Wolfe et al. 2005). In the current implementation of this paradigm, speakers produced conceptually ambiguous texts and listeners subsequently listened to these texts preceded by a context that in some cases did (highly informative or intermediately informative title) and in others did not (no title) facilitate the extraction of a coherent situation model. The informative context renders available an appropriate event schema that can guide the interpretation of incoming information over time. Schemas are templates that are retrieved when new information needs to be interpreted (van Kesteren et al. 2012; Gilboa and Marlatte 2017). On the neural level, cognitive schemas are represented as associative networks that influence processing of new information. In the present study, an appropriate event schema to guide the construction of a coherent situation model was only provided when the title was highly informative, and to a limited degree when the title was intermediately informative. By keeping all bottom-up sensory and linguistic information identical, this manipulation allowed us to isolate behavioral (comprehension, recall) and hemodynamic effects that were related to situation model processing.

Moreover, in order to shed light on the neural underpinnings that reflect a shared situation model representation across speakers and listeners, we investigated the similarity of the BOLD time course across participants, using an inter-subject correlation (ISC) approach (Hasson et al. 2004). The rationale of this approach is that processing similar information (e.g., the same situation model) across individuals should be reflected by similar hemodynamic response time courses in the neural regions involved in processing relevant information. Here, ISC was calculated between the speakers’ and listeners’ hemodynamic time courses. Previous research provided insight into the relationship of the neural time course between speakers and listeners of a narrative, using an ISC approach (Stephens et al. 2010). The previous approach, however, could not disentangle which part of the ISC reflected the shared situation model and which part reflected shared linguistic and sensory processing. The current study was designed to provide a clearer picture of the neural basis of shared representations across interlocutors related to the situation model. In summary, by using this approach of combining the ambiguous text paradigm, conventional BOLD activation and ISC analyses, we aimed at (1) disentangling the neurocognitive processes underlying the construction of a situation model in multi-sentence discourse and at (2) providing new insight into the relationship of the neural activity between speakers and listeners when they share rich and complex conceptual representations, such as situation models.

### 1.1. Hypotheses

If supportive contextual information, i.e., a highly or intermediately informative title, is provided before the ambiguous text, it can activate an appropriate event schema that helps interpreting ambiguous passages (Bransford and Johnson 1972). The contextual information reduces the uncertainty about the discourse and, on the behavioral level, is expected to improve understanding and recall of its content (Bransford and Johnson 1972; Wahlberg and Magliano 2004; Ames et al. 2015). Thus, the behavioral scores for comprehension and recall were expected to be highest when a highly informative title (HT) was given, followed by the intermediately informative title condition (IT) and least when no title (NT) was given. On the neural level, the regions where we expected to find increased BOLD activation in listeners and increased ISC between speakers and listeners when sharing a situation model, were (1) the ventromedial prefrontal cortex (vmPFC), (2) the posterior cingulate cortex (PCC) and precuneus, and (3) the right inferior frontal gyrus (rIFG).

To conclude, in the highly informative title condition (HT), the behavioral scores for comprehension and recall were expected to be highest, and the hemodynamic measures (BOLD, ISC) related to situation model processing to be strongest in the regions mentioned above associated with (large-scale) conceptual processing. In the intermediately informative title condition (IT), comprehension and recall as well as the hemodynamic measures were expected to be reduced, yet superior to the no title condition (NT).

## 2. Materials and Methods

### 2.1. Participants

Participants were 42 right-handed native speakers of Dutch (31 female, 11 male) between 18 and 35 years of age (average age: 22.6 ± 3.2 (SD) years), with no history of neurological, psychiatric or language disorders. Fifteen participants (12 female, 3 male) acted as *speakers*, and 27 participants (19 female, 8 male) as *listeners*. All participants had normal audition and normal or corrected-to-normal vision. The data of three participants (one speaker, two listeners) were excluded from analysis due to excessive head motion during the main task. All participants gave written informed consent and received payment or course credit. The study was approved by the local ethics committee (*Commissie Mensgebonden Onderzoek regio Arnhem-Nijmegen*).

### 2.2. Stimuli and experimental design

The *speakers* were asked to read aloud short expository texts in Dutch that contained a topic description (short familiar event scripts; e.g., *to make coffee*), which were later presented to the *listeners.* Here we used ambiguous expository texts, i.e., paragraphs that are globally coherent only when an informative title that describes the topic is provided in the beginning (see also, Dooling and Lachman 1971; Bransford and Johnson 1972; Wahlberg and Magliano 2004). This is because when no title is given, a global coherence of the text and the comprehension of the topic are difficult to establish. By using these texts, our aim was to manipulate the depth of conceptual processing via the informativeness of the contextual cue (title). Fifteen texts, each describing one of the following topics, were produced by each of the participants in the speaker’s group: e.g., *to make coffee, to wash clothes, to carve a pumpkin, horseback riding, to build a snowman, to wash dishes, to water flowers* (for an example of a topic description, see Table 1; for a full list of topics and tiles, see Supplementary Information, Table S 1). Some topic descriptions were adapted from previous studies (Smirnov et al. 2014; Ames et al. 2015) and others were newly created. All topics involved scripts, i.e., daily routines of human behavior, or descriptions of activities that were comparable with respect to self-relatedness. Maintaining a similar degree of self-relatedness across texts was considered relevant due to the role of some critical functional regions, e.g., dmPFC, in self-referential processes (see for instance, Gusnard et al. 2001; Northoff and Bermpohl 2004; Sieborger et al. 2007).

**Table 1.** Example of a topic description and corresponding titles (translated from Dutch).

| **Title** | **Highly informative** | **Intermediately informative** | **No title** |
| --- | --- | --- | --- |
|  | *To make coffee* | *To prepare a beverage* | *#####* |
| **Text** | *This process is often full of rituals. It happens at all times of the day. The process also occurs everywhere, and sometimes there are many people involved while at other times it is carried out alone. The process does not take a lot of time, but requires a number of accessories. In general, it is done frequently, but the quantity differs depending on the producer. In addition, there is also no agreement on the best way how to do it. There are several ways how to proceed, which lead to different outcomes. Producers often have different preferences, and often find it easy to distinguish the quality of the product. Sometimes the final product is adapted by the producer according to individual preferences. It happens everywhere in the world and in almost every culture, even though the process can be very different. The positive and negative aspects of the process are a matter of discussion, but more and more consensus seems to come about.* |  |  |

For each topic, two titles were identified that varied in their degree of *Title informativeness* about the topic, i.e., (1) highly informative (e.g., *To make coffee*), and (2) intermediately informative (e.g., *To prepare a beverage*). The titles for all 15 texts can be found in Supplementary Information, Table S 1. In an additional condition (3) no title but only a series of hashmarks was presented (e.g., *#############*). Speakers were presented with only the highly informative title before the text production, whereas listeners were presented with one of the three *Title informativeness* conditions for each text. In order to evaluate the ambiguity of the texts and the influence of the titles on text comprehension, a pretest was conducted on an independent sample of participants. Details on the pretest are provided in Supplementary Information, Methods, Pretest.

### 2.3. Task and procedure

The experiment was presented using Presentation software (Neurobehavioral Systems; version 19.0). Participants performed either the speaking (*speakers*) or listening (*listeners*) task while lying in the MRI scanner.

#### Production task

Before entering the scanner, *speakers* were given time to familiarize with all 15 ambiguous texts, preceded by the disambiguating highly informative title. They were instructed to read each text on their own pace so as to get to know the topic and to be able to afterwards read the texts aloud fluently in the scanner. In the scanner, each trial started with the presentation of a green fixation cross for 4 s, followed by the visual presentation of the highly informative title in the center of the screen for 4 s (Figure 1). Then, a white fixation cross was presented for the remaining 6 s preceding the text, followed by the visual presentation of the entire text on the screen. Participants were instructed that in each trial, the presentation of the text on the screen indicated the onset for the production of the expository text which would remain on the screen until the speaker indicated the end of the production with a button press. Next, the white fixation cross was again visible for 6 s after the offset of the text production. In total, each speaker produced 15 texts (mean duration of text production: 61.5 ± 7.1 (SD) sec), which were presented in a constant order across speakers. The order was maintained constant in order to account for any conceptual influence between texts presented in sequence. All 15 texts were produced in one run, from which three texts per speaker were subsequently chosen, according to the quality of the recording. Recordings from 5 different speakers were subsequently presented to 9 different listeners. In total, 3 groups representing this combination between speakers and listeners were formed.

**Figure 1.**
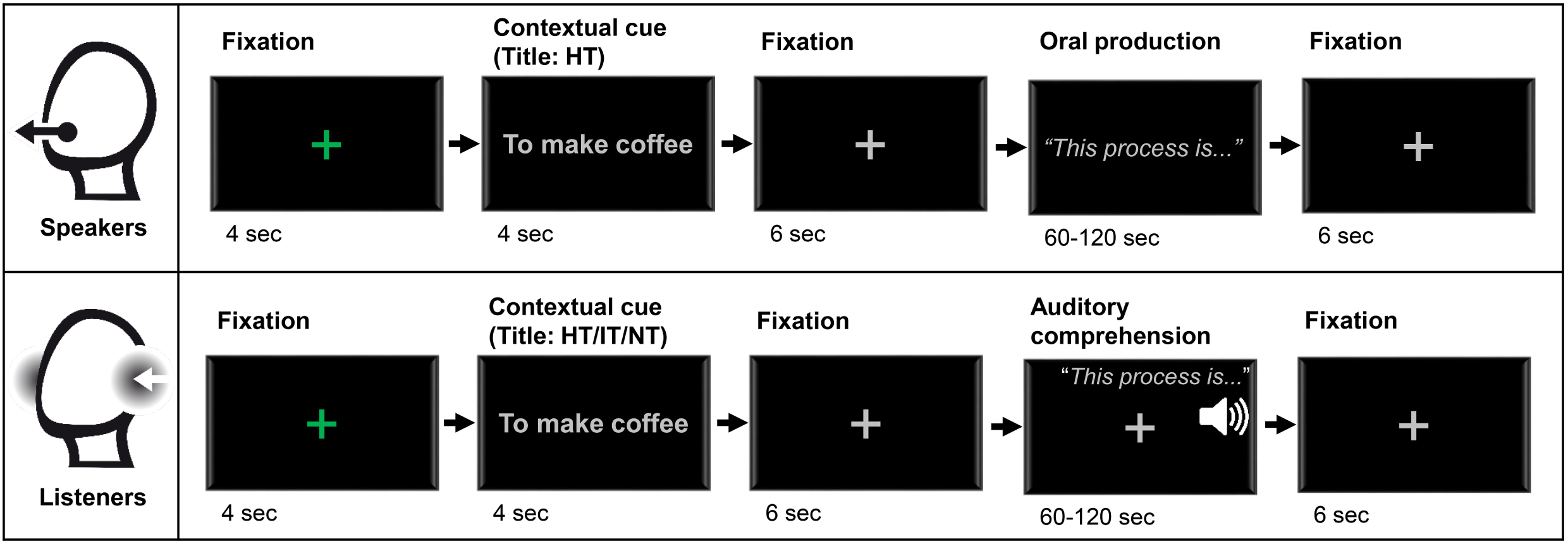
Time course of a trial for speakers and listeners. HT, Highly informative title; IT, Intermediately informative title; NT, No title.

#### Comprehension task

The *listeners* listened to the expository texts while lying in the MRI scanner. They were instructed to listen to and to try to understand each of the texts as well as possible, and to indicate with a button press when they believed to have understood the topic of the text during listening. Before entering the scanner, they were presented one training trial in order to familiarize them with the task. In total, 15 texts were presented to each listener in a constant order across listeners. The order of texts was identical for speakers (production task) and listeners (comprehension task). The assignment of auditory texts to the experimental conditions (highly informative/intermediately informative/no title) was counterbalanced across participants, i.e., each listener was presented each topic description only once, preceded by one of the titles or the hashmarks. The 15 texts were assigned to one of the three conditions in a pseudorandomized order (max. three texts per condition in immediate succession). Counterbalancing ensured that across listeners, all texts were presented an equal number of times in each condition. Each listener was presented 5 texts per condition and 3 per speaker. Each trial started with the presentation of a green fixation cross for 4 s, followed by the visual presentation of the title in the center of the screen during 4 s (Figure 1). Then a white fixation cross was presented during the 6 s preceding the auditory presentation of the text and stayed on screen until 6 s after the offset of the text. The expository texts were presented in one run consisting of 15 texts.

#### 2.3.1. Comprehension and recall measurements

After scanning, the listeners were asked to carry out a comprehension and a recall task for each expository text (see also, Bransford and Johnson 1972; Smirnov et al. 2014; Ames et al. 2015). To do so, all texts were presented auditorily a second time, in the same order as previously in the scanner. Following the presentation of each title and text, the participants were asked to evaluate how easy it was to understand the text using a scale ranging from 1 (“I was totally confused”) to 7 (“It was all totally clear”). Then they were asked to identify and write down the topic of the text and to indicate their confidence in this answer on a scale from 1 (“I am guessing randomly”) to 7 (“I am totally certain”). This comprehension task was followed by an open recall task in which participants were asked to recall as many ideas as possible from the text and to write them down. For this open recall task, the “idea units” had been identified a priori and corresponded to either individual sentences, basic semantic propositions, or phrases related to the topic. The texts contained on average 27.4 ± 3.9 (SD) idea units (range 21-35). The recall performance was scored using the list of idea units and was subsequently rescaled as the ratio of recalled idea units to maximum number of possible idea units per text.

#### 2.3.2. Auditory recording and auditory stimulus presentation

Participants were equipped with in-ear phones (Sensimetrics), which also provided ear-protection, and for speakers MRI-compatible microphones were also provided (Optical Microphone FOMRI™ III Dual Channel Microphone System for fMRI). These devices use built-in digital signal processing algorithms to reduce acoustic scanner noise from the audio presentation and/or the recording, respectively. The speakers’ recorded audio files were subsequently cleaned from residual scanner noise using Adobe Audition CS6 (version 5.0.2). This cleaning procedure involved first the capturing of a noise print in the initial part of each sound file figuring only scanner noise before speech onset, which was then filtered out from the entire sound file (reduction by 40 dB, FFT size: 2048). Finally, all sound files were intensity normalized (target intensity: 65 dB) in Praat (version 6.0.36; Boersma and Weenink 2016). In the scanner, before the first functional scan was obtained, a sound check was carried out in which segments of stimuli from the main task were presented in order to individually adjust the stimulation volume to be heard over the scanner noise.

#### 2.3.3. Manual response

Participants were equipped with a button box at the right hand (Curved Lines HHSC-2x4-C; fORP 932 Response Box Interface).

### 2.4. Localizer tasks

After the main task, three functional localizer tasks were run in the following order: a *False belief localizer* (Dodell-Feder et al. 2011) in order to target the core theory of mind (ToM) network, a *WM/DMN (working memory/default mode network) localizer* and a *Language localizer* (shortened version of Lam et al. 2016; Schoffelen et al. 2019). Details on the localizer tasks are provided in Supplementary Information, Methods, Localizer tasks.

### 2.5. fMRI data acquisition and preprocessing

The participants’ neural activity was recorded in a Siemens 3T MAGNETOM Prisma MRI scanner using a 32-channel head coil. Functional images with 2 mm isotropic resolution were acquired using a T2*-weighted echo-planar imaging sequence (TR: 1000 ms, TE: 34 ms, 66 axial slices per volume, FOV = 210 mm, 60° flip angle, interleaved multi-slice mode, multi-band acceleration factor: 6). Moreover, two fieldmaps (TR: 620 ms; TE1: 4.70 ms, TE2: 7.16 ms) were acquired, one after the main task and a second one after the three localizer tasks. In addition, a T1-weighted anatomical scan with 1 mm isotropic resolution (TR: 2300 ms, TE: 3.03 ms, 192 sagittal slices, FOV = 256 mm, 8° flip angle) was acquired for each participant. During the functional scans, the left eye was tracked using a non-invasive, video-based eye tracking system (SMI-eyetracker; iView X) in order to assure that the participants stayed awake in the scanner.

The functional data were preprocessed and analyzed using FSL (FMRIB Software Library, version 5.0.11; www.fmrib.ox.ac.uk/fsl; Jenkinson et al. 2012). The preprocessing included: deletion of first 5 volumes, brain extraction (FSL’s BET; Smith 2002), motion correction (FSL’s McFLIRT), correction of geometric distortions due to magnetic field inhomogeneities using a fieldmap (FSL’s FUGUE), spatial smoothing with a Gaussian 5 mm FWHM kernel, and temporal high-pass filtering at 100 s. The image registration was performed in two stages involving, first, the linear registration of the skull-stripped functional images to the skull-stripped high resolution T1-weighted structural images (FSL’s Linear Registration Tool (FLIRT); Jenkinson and Smith 2001; Jenkinson et al. 2002), followed by the nonlinear registration to the standard space MNI152 (Montreal Neurological Institute) 2 mm brain template (FSL’s NonLinear Registration Tool (FNIRT); Andersson et al. 2007) (12 degrees of freedom, full search, 10 mm warp resolution).

#### 2.5.1. Physiological noise correction

Physiological data were acquired (sampling rate: 5000 Hz; BrainVision Recorder, Brain Products, Gilching, Germany) using a respiratory belt and a heart rate monitor (pulse oximeter on a finger). For two participants only the respiratory trace was included due to poor quality of cardiac measures. FSL’s PNM (Brooks et al. 2008) was used to convert the respiratory and cardiac traces into 14 physiological regressors. These voxelwise confound regressors were added in the conventional BOLD GLM analysis. In the inter-subject correlation (ISC) analysis, the voxelwise confounds were regressed out from the preprocessed functional data before splitting the functional data into segments corresponding to the individual stimuli, and subsequent ISC analysis.

### 2.6. Statistical analyses

#### 2.6.1. Behavioral data analyses

Comprehension and recall measures obtained from the listeners after scanning were analyzed in a mixed-effects model, including the fixed effect *Title informativeness* (high(HT)/intermediate(IT)/none(NT)), as well as the intercepts for the random effects *Subject* and *Item (Text)*, and the by-*Subject* and by-*Item* random slopes for the effect of *Title informativeness*. The linear mixed-effects analysis was carried out using the *lmer()* function from the *lme4* package (Bates et al. 2015), and the optimizer method *L-BFGS-B* from package *optimx* (Nash and Varadhan 2011) as well as the *anova()* function from the *stats* package. Degrees of freedom were approximated using *Satterthwaite*’s method. Post-hoc tests were carried out using a Tukey-test. Bonferroni adjustment of *p*-values was done via the *glht()* function from the *multcomp* package (Hothorn et al. 2008) in R version 3.5.1 (R Core Team 2014).

#### 2.6.2. fMRI data: Conventional BOLD activation analyses

For the group-level analysis, individual participant first-level models were created using a general linear model (GLM) with *Title informativeness* condition (high(HT)/intermediate(IT)/none(NT)) as explanatory variable of interest and gamma convolution of the hemodynamic response function. Individual texts were treated as blocks of stimulation in our design. Six standard motion parameters as well as physiological noise regressors issued from FSL’s PNM (see section 2.5.1) were included in the statistical model as voxelwise confound regressors. In the individual participant first-level analysis, whole-brain *z*-maps were family-wise error (FWE)-corrected at cluster-level *p*<.05 (*z*=2.3). Higher-level random effects analysis using FSL’s FLAME (FMRIB’s Local Analysis of Mixed Effects) was performed on the contrast images obtained from the first-level analysis. In the group level analysis, whole-brain *z*-maps were FWE-corrected at cluster-level *p*<.05 (*z*=3.1). The cluster maximum is reported for each cluster with its respective *z*-value, as well as further local maxima if they fell within distinct anatomical regions. All reported coordinates are in MNI space and anatomical labels were attributed according to the *Harvard-Oxford cortical* and *subcortical structural atlases* and the *Cerebellar Atlas in MNI152 space after normalization with FNIRT*. As for differential contrasts, to further explore the underlying activation pattern driving any significant effects, the voxel with the maximal *z*-score of each significant cluster was identified, and the average percent signal change for this voxel for each simple positive contrast (HT, IT and NT respectively vs baseline) was extracted and plotted in a bar-graph for the three conditions of *Title informativeness*. Finally, in addition to the whole-brain analyses, region of interest (ROI) analyses were conducted. Our functional regions of interest (fROIs) were based on contrasts obtained from the localizer tasks (for details, see Supplementary Information, Methods, Localizer tasks). Within each fROI, the percent signal change for each *Title informativeness* condition (simple positive contrast vs baseline) was extracted for each participant. A two-way repeated measures ANOVA including the fixed effects *Title informativeness* and *ROI* was run using the function *aov()*. Only a main effect of *Title informativeness* or a *Title informativeness* by *ROI* interaction was further inspected using the function *emmeans()*.

#### 2.6.3. fMRI data: Inter-subject correlation (ISC) analyses

An inter-subject correlation (ISC) approach (Hasson et al. 2004) was used to assess the similarity of the stimulus-related time course of neural activity across individuals. For stimuli with an extended time course and a temporally varying and complex BOLD response, an inter-subject correlation (ISC) approach can be more sensitive than conventional GLM analyses, because ISC analyses primarily assesses the comparability of the BOLD response pattern over time across participants without requiring a model of the BOLD response at specific time points (Smirnov et al. 2014; Ames et al. 2015). In ISC analysis, the correlation between each voxel’s time course in one participant and the time course of the equivalent voxel in a second participant’s brain is calculated. Brain areas that manifest a high correlation between the two participants’ time courses are considered to play a role in processing stimulus-specific information that is common across participants.

Before calculating ISC maps, the six standard motion parameters as well as the voxelwise confounds (physiological noise; cf. section 2.5.1) were regressed out from the preprocessed functional data. The preprocessed and cleaned functional data in MNI space were then split into stimulus-specific segments, excluding the first 12 volumes (= 12 seconds) in order to deal with stimulation-initial hemodynamic transients (Boynton et al. 1996; Logothetis et al. 2001).

Next, the correlation maps for each stimulus and speaker-listener pair were calculated using in-house customized code in Matlab. Moreover, ISC maps for each speaker-listener pair, between unmatched stimuli were calculated to obtain a baseline condition. All pairwise correlation maps were then normalized (Fisher’s r-to-z transform) and averaged per speaker-listener pair and condition. Then, in order to account for spatio-temporal dependencies in the data, the averaged ISC maps per speaker-listener pair and condition were analyzed with a (Wilks’ λ) permutation test (1.000 permutations), using FSL’s PALM (Permutation Analysis of Linear Models; Winkler et al. 2014). Repeated measures were accounted for by including mean effect regressors for each participant pair and by allowing permutation exchangeability only between samples from the same participant pair across conditions. The whole-brain analysis was limited to only gray matter voxels by using an average gray matter mask obtained from all participants’ structural scans. Whole-brain *z*-maps were FWE-corrected at cluster-level *p*<.05 (*z*=2.3). All reported coordinates are in MNI space and anatomical labels were attributed according to the *Harvard-Oxford cortical structural atlas* and the *Brodmann atlas*.

In addition to the whole-brain analyses, ROI analyses were conducted for the ISCs. Our fROIs were based on contrasts obtained from the localizer tasks (for details, see Supplementary Information, Methods, Localizer tasks). Within each ROI, mean ISC values were extracted for each participant pair and condition. A two-way repeated measures ANOVA including the fixed effects *Title informativeness* and *ROI* was run using the function *aov()* in R for each ISC combination. Only a main effect of *Title informativeness* or a *Title informativeness* by *ROI* interaction was further inspected using the function *emmeans()*.

Finally, we wanted to take into account the relationship between ISCs and behavioral measures of text comprehension and recall. Average speaker-listener ISCs were calculated per text and condition. Correlation analyses between these average ISC values and average behavioral scores (comprehension, recall) per text and condition were conducted. These tests were carried out using a (Wilks’ λ) permutation test (1.000 permutations) in FSL’s PALM (Winkler et al. 2014). Data and design were demeaned and repeated measures were accounted for by including mean effect regressors for each text and by allowing permutation exchangeability only within each text across conditions. Moreover, ROI analyses were conducted using the *rmcorr()* function from the *rmcorr* package (Bakdash and Marusich 2017) in R, to calculate the repeated measures correlation (rmcorr) coefficient per ROI. The calculation of the rmcorr coefficient takes into account the corresponding repeated measures per stimulus text across conditions when analyzing a correlation.

## 3. Results

### 3.1. Behavioral data

Comprehension scores and recall scores were compared across three different levels of *Title informativeness* (HT/IT/NT) using a mixed-effects model. Behavioral results are presented in Figure 2. Comprehension scores significantly differed between conditions (*F*(2,18)=48.75, *p*<.001), and post-hoc analyses revealed that the ease of comprehension was rated significantly higher in the HT (5.46 ± 0.21 (SE), 95% CI[5.06, 5.87]) than the IT (4.75 ± 0.30 (SE), 95% CI[4.16, 5.34]; *z*=2.86, *p*_adj_<.05) and NT (3.25 ± 0.21 (SE), 95% CI[2.84, 3.65]; *z*=9.74, *p*_adj_<.001) conditions, and in the IT than the NT (*z*=4.76, *p*_adj_<.001) condition. Similarly, the recall performance differed between conditions (*F*(2,19)=7.11, *p*<.01). Post-hoc comparisons revealed that significantly more idea units were recalled in the HT (21.40 ± 2.17 (SE) %, 95% CI[17.13, 25.66]) than the NT (15.91 ± 1.65 (SE) %, 95% CI[12.66, 19.15]; *z*=3.76, *p*_adj_<.001) condition, and in the IT (20.08 ± 1.98 (SE) %, 95% CI[16.19, 23.98]) than the NT (*z*=2.56, *p*_adj_<.05) condition. Hence, text comprehension and the recall of ideas from each text were strongly dependent on the information provided in the title, with highly informative titles leading to best comprehension and recall scores, no titles to worst scores, and intermediately informative titles to in-between performance.

**Figure 2.**
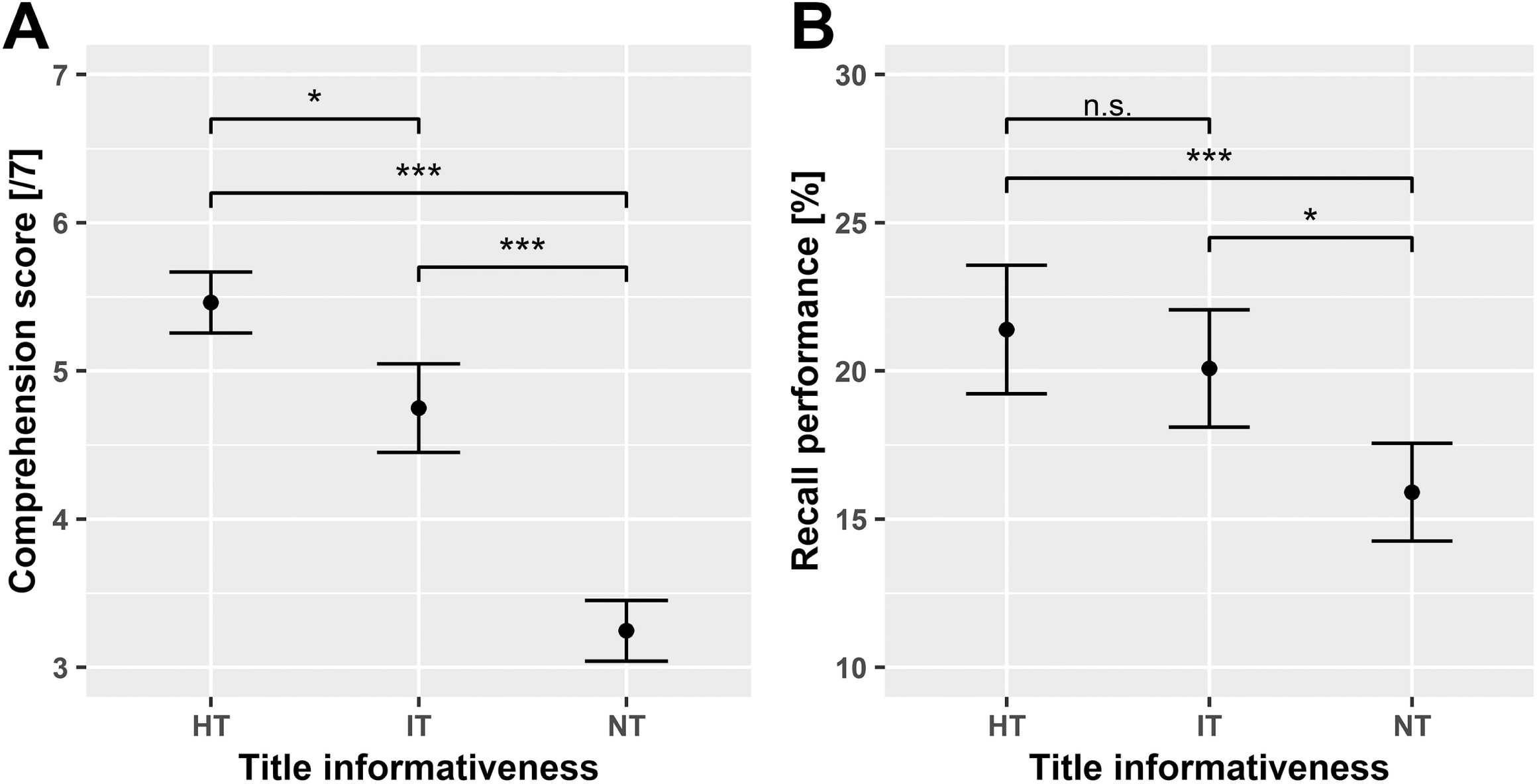
Post-scanning comprehension (A) and recall (B) measures in listeners. Error bars represent ±1 SD. HT, Highly informative title; IT, Intermediately informative title; NT, No title. ***, *p*<.001; **, *p*<.01; *, *p*<.05; n.s., not significant; Bonferroni-adjusted.

### 3.2. fMRI data: Localizer tasks

Functional regions of interest (fROIs) selected with the localizer tasks were the following (see also Supplementary Information, Table S 2 and Figure S 1): The selected *theory of mind (ToM)* fROIs lay in the right medial prefrontal cortex (center of the ROI at MNI coordinate [6/58/28]), the left temporal pole [-56/4/-22], the (right) medial parietal cortex (precuneus [2/-56/36]) and in the right inferior lateral parietal cortex (angular gyrus [58/-54/30]). The selected *working memory* fROIs were in the right insula [36/20/-2], right superior lateral parietal lobule (angular gyrus [38/-54/44]) and in the right middle/superior frontal gyrus [28/6/56]. The selected *default mode* fROIs lay in the left medial prefrontal cortex [-6/62/22], right hippocampus [26/-16/-16] and in the left medial parietal cortex (posterior cingulate [-4/-50/24]). The selected *language processing* fROIs were in the left middle/superior temporal gyrus [-56/-8/-12] and in the left inferior frontal gyrus [-54/20/22].

### 3.3. fMRI data: Conventional BOLD activation analysis

To see which regions were more active when processing conceptually coherent compared to incoherent discourse, a conventional GLM analysis was run on the listeners’ hemodynamic response, with levels of *Title informativeness* (HT, IT, NT) as the explanatory variable of interest. Each simple positive contrast, i.e., HT, IT, and NT, vs baseline, showed a strong involvement of mostly left-lateralized fronto-temporal regions, including the superior temporal cortex bilaterally and the inferior frontal gyrus, partes orbitalis, triangularis and opercularis in the left hemisphere and partes orbitalis and triangularis in the right hemisphere (threshold *z*=3.1, *p*<.05 at cluster-level; for simple contrast maps, see Figure 3A). Furthermore, in all three conditions, significant BOLD activation was found in the left dorsomedial prefrontal cortex (dmPFC) and supplementary motor area (SMA), in the right precentral gyrus and in the cerebellum. However, only in the HT and IT conditions, significant BOLD activation was found in the left ventromedial prefrontal cortex (vmPFC).

**Figure 3.**
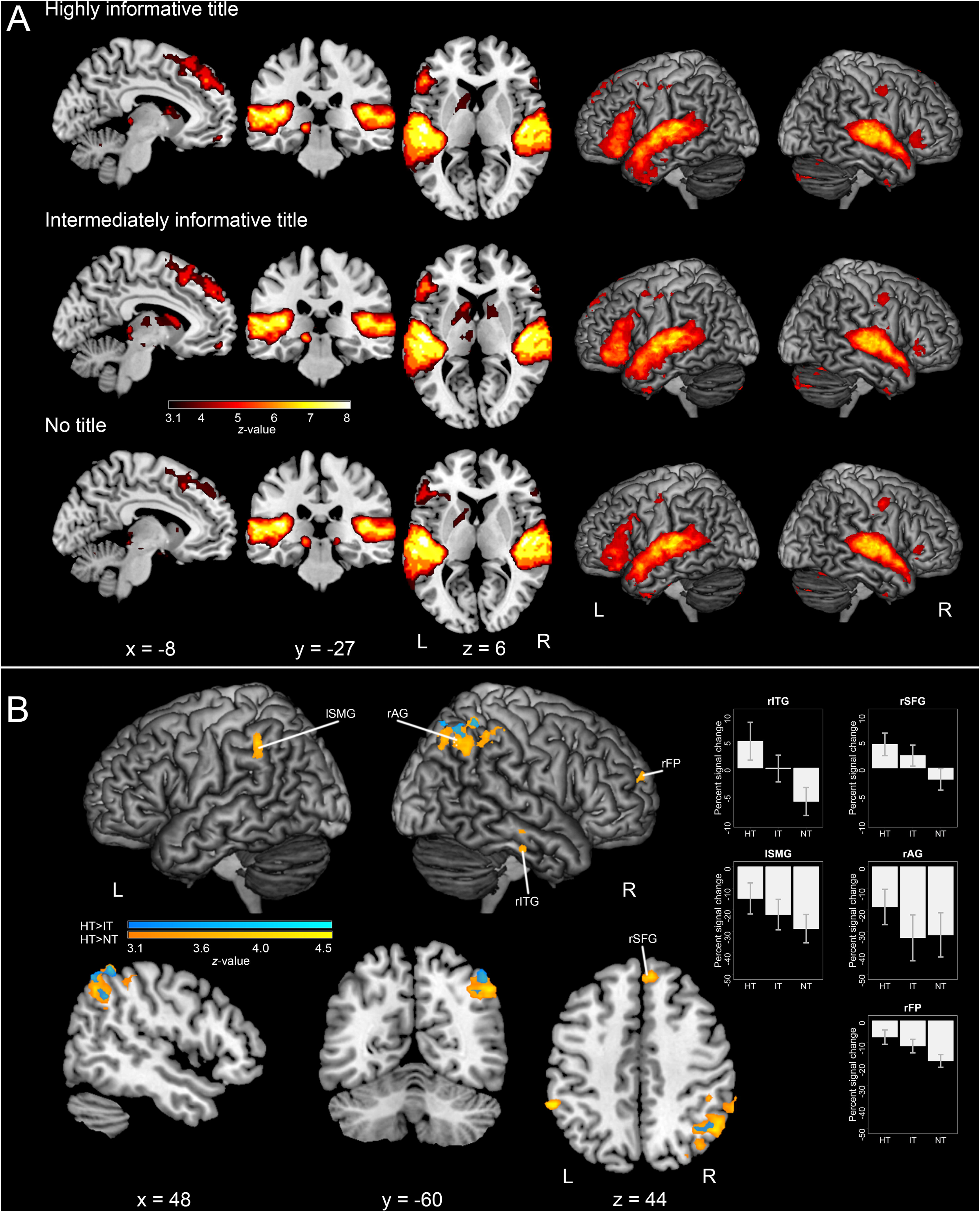
Conventional BOLD activation analysis in listeners. **A** Significant BOLD activation in the positive contrasts for each of the three conditions (highly informative title/HT, intermediately informative title/IT, no title/NT) is displayed. Whole-brain *z*-map, FWE-corrected at cluster-level *p*<.05 (*z*=3.1). **B** Clusters showing a significant BOLD activation effect for the contrasts HT > IT (cool colors) and HT > NT (warm colors) are displayed. Whole-brain *z*-map, FWE-corrected at cluster-level *p*<.05 (*z*=3.1). Bar-graphs represent the percent signal change for the peak voxel in each cluster (contrast HT > NT). Error bars represent ±1 SEM. HT, Highly informative title; IT, Intermediately informative title; NT, No title. lSMG, left supramarginal gyrus; rAG, right angular gyrus; rFP, right frontal pole; rITG, right inferior temporal gyrus; rSFG, right medial superior frontal gyrus / dorsomedial prefrontal cortex.

Differential contrasts revealed that the right inferior lateral parietal cortex, notably the angular gyrus (AG), showed a higher BOLD response in the HT compared to the IT and NT conditions, a pattern that was similar in the left supramarginal gyrus where the HT showed a higher BOLD response compared to the NT condition (Figure 3B and Table 2). Moreover, the right frontal pole showed a higher BOLD response in the HT and IT compared to the NT condition. Finally, the right superior frontal and paracingulate gyri (medial frontal cortex) as well as the right posterior inferior and middle temporal gyri revealed stronger activation in the HT compared to the NT condition.

**Table 2.** MNI coordinates of peak activations obtained in the whole-brain GLM analyses.

| Contrast | Cluster | Hemisph. | Region name | Z | Peak MNI |  |  |
| --- | --- | --- | --- | --- | --- | --- | --- |
|  | extent |  |  | value | coordinate |  |  |
|  | (voxels) |  |  |  | x | y | z |
| HT>IT | 188 | R | superior lateral occipital cortex | 4.01 | 34 | -70 | 58 |
|  |  | R | angular gyrus | 3.88 | 40 | -46 | 34 |
| HT>NT | 823 | R | angular gyrus | 4.43 | 48 | -50 | 60 |
|  |  | R | superior lateral occipital cortex | 4.11 | 46 | -60 | 46 |
|  | 113 | L | supramarginal gyrus | 4.45 | -58 | -42 | 44 |
|  | 106 | R | frontal pole / prefrontal cortex | 3.9 | 22 | 56 | 18 |
|  | 97 | R | superior frontal gyrus / dorsomedial prefrontal cortex | 4.34 | 10 | 42 | 40 |
|  |  | R | paracingulate gyrus | 3.35 | 8 | 34 | 38 |
|  | 94 | R | posterior inferior/middle temporal gyrus | 3.93 | 60 | -18 | -28 |
| IT>NT | 155 | R | frontal pole / prefrontal cortex | 4.13 | 22 | 58 | 20 |
| NT>HT | 127 | R | vermis VIII | 4.15 | 6 | -60 | -34 |
|  |  | R | cerebellum I-IV | 4.14 | 6 | -46 | -26 |
| IT>HT | - |  |  |  |  |  |  |
| NT>IT | - |  |  |  |  |  |  |

Whole-brain $z$ -maps were FWE-corrected at cluster-level $p<.05$ ( $z=3.1$ ). Hemisph., Cerebral hemisphere; HT, Highly informative title; IT, Intermediately informative title; NT, No title.

In the ROI analysis, the two-way repeated measures ANOVA revealed a *Title informativeness* by *ROI* interaction (*F*(22,528)=1.90, MSE=115, *p*<.01). Post-hoc analyses showed that the BOLD response was significantly higher in the HT compared to the NT condition in the (right) precuneus, and in the right inferior and superior lateral parietal cortex (angular gyrus) (*p*s<.01; Tukey-adjusted).

To sum up, all three *Title informativeness* conditions (HT, IT, NT) were associated with strong activation in the left inferior frontal cortex - and to a smaller extent in the right inferior frontal cortex, and in bilateral superior temporal regions, which are typically associated with (auditory) language processing. Differential contrasts revealed that the bilateral lateral parietal cortex (supramarginal gyrus, angular gyrus) and the medial parietal cortex (precuneus) showed a stronger BOLD response in the highly informative than the no-title condition, with the intermediately informative condition showing values in-between. In the right hemisphere, clusters in the medial and lateral prefrontal cortex as well as in the inferior temporal cortex showed a similar pattern.

### 3.4. fMRI data: Speaker-listener ISC analysis

For speaker-listener ISCs, the whole-brain analysis showed that ISCs were stronger in the right dorsolateral prefrontal cortex in the IT condition compared to the unmatched-text baseline (marginally significant; *p_corr_*<.10). No other effects were found in the whole-brain or ROI analyses.

The whole-brain correlation analysis with behavioral measures revealed a positive correlation between the listeners’ comprehension scores and speaker-listener ISCs in the bilateral (ventro)medial prefrontal cortex (vmPFC; *p_corr_*<.05; Figure 4A). The ROI correlation analysis additionally indicated a positive correlation between the listeners’ recall performance and ISCs in two regions, namely the (right) precuneus (*p_uncorr_*<.05) and the right hippocampus (*p_uncorr_*<.05; Table 3).

**Figure 4.**
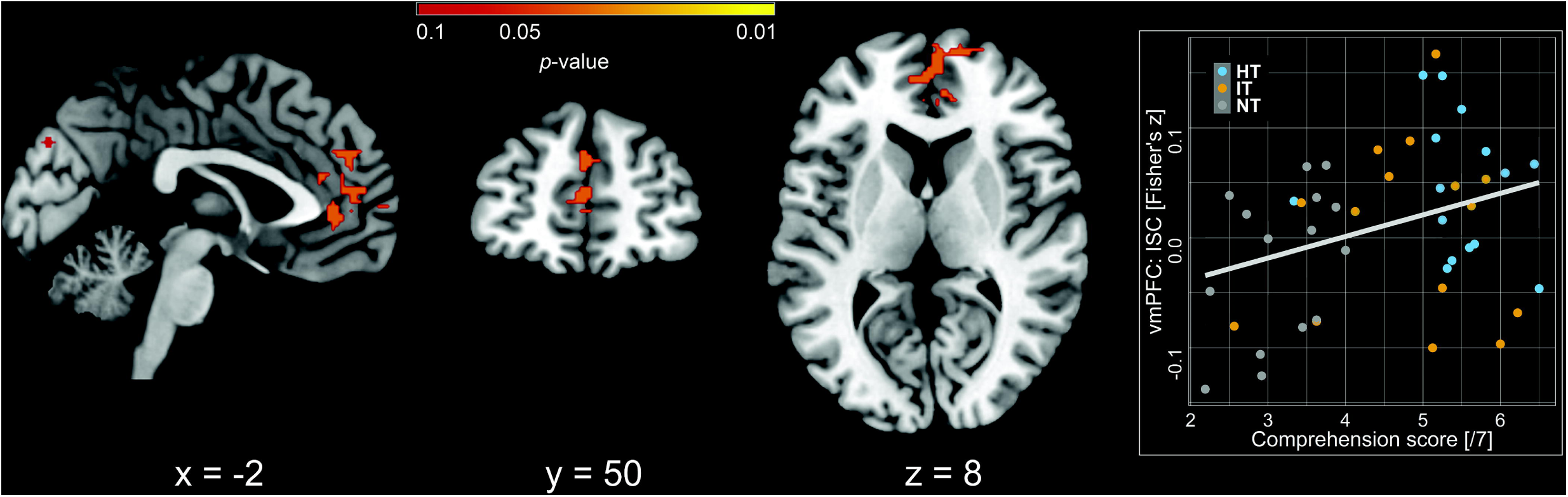
Inter-subject correlations (ISC) correlated to text comprehension scores. Clusters with significant correlations between behavioral comprehension measures and speaker-listener ISCs (Fisher’s *z* transform of Pearson’s correlation coefficient) are displayed. Correlations were conducted for average comprehension scores and ISCs across all three conditions (HT, IT, NT). The correlation plot represents ISCs for voxel -3/52/12 (left ventromedial prefrontal cortex; vmPFC) Whole-brain significance (*p*-value) map, FWE-corrected at cluster-level *p*<.05 (*z*=2.3).

**Table 3.**
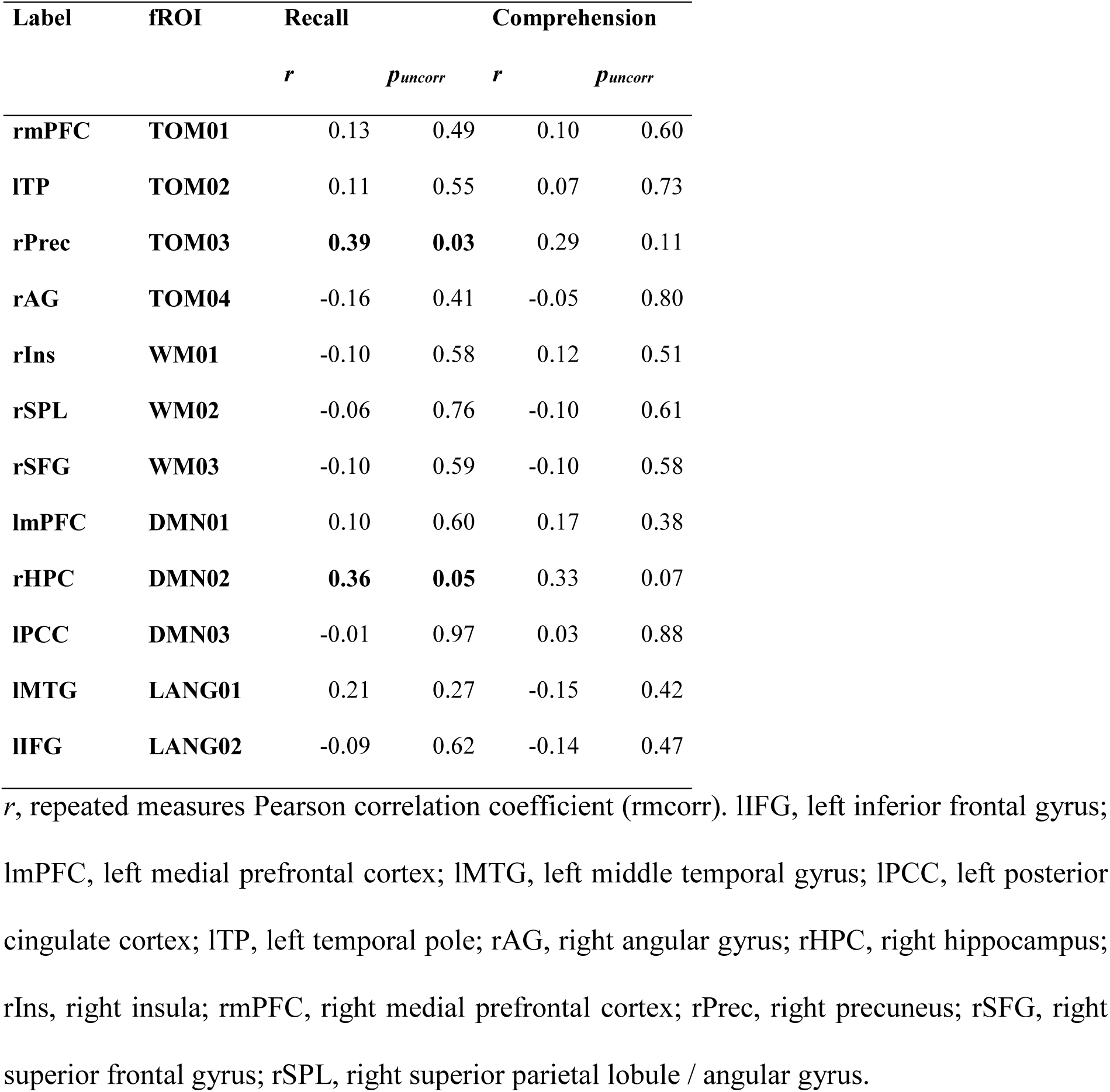
ROI analysis on the correlation between inter-subject correlations and behavioral measures. Repeated measures correlations between speaker-listener inter-subject correlations (ISC) and behavioral measures (comprehension, recall) in the 12 fROIs targeting core theory of mind (TOM), default mode (DMN), verbal working memory (WM) and language processing regions are displayed. Uncorrected, significant correlations are highlighted in bold font.

To sum up, the ISC analyses revealed that speakers’ and listeners’ BOLD time courses were more aligned in the medial prefrontal cortex when the listeners subsequently reported a good text comprehension. Similarly, the speaker-listener alignment tended to be higher in the precuneus and hippocampus when the listener’s subsequent recall of text elements was good.

## 4. Discussion

In the present study, our first main goal was to disentangle the neurocognitive processes underlying the construction of a situation model in multi-sentence discourse. Moreover, our second aim was to identify the neural bases reflecting shared representations in speakers and listeners of discourse. To investigate these questions, we implemented a variant of the ambiguous text paradigm (Bransford and Johnson 1972). In the MRI scanner, speakers produced ambiguous texts, preceded by a highly informative title that allowed them to produce the texts within a coherent situation model. Subsequently, listeners listened to these texts, preceded by either a highly informative title (HT), an intermediately informative title (IT) or no title at all (NT). In so doing, we aimed at disentangling the neurocognitive processes involved in discourse processing and at identifying the shared neurocognitive processes between speakers and listeners who successfully shared a situation model. We expected situation model construction to be most elaborate when a highly informative title was given (HT), and to a minor degree when the title was intermediately informative (IT), because the title allowed to activate a cognitive schema to guide the comprehension of incoming information in the text. In general, discourse processing was found to be associated with hemodynamic activation in the classical left-dominant fronto-temporal language network, but only the successful processing of a coherent discourse level conceptual representation (situation model) additionally involved activation in the bilateral lateral parietal, medial parietal and prefrontal regions, as well as the hippocampus. More specifically, across listeners, when they were able to establish a situation model similar to that of the speaker, (shared) activation was observed in the bilateral lateral parietal regions (angular gyrus, supramarginal gyrus), as well as in the medial prefrontal cortex. Moreover, good recall of text elements in listeners was correlated with speaker-listener alignment of the BOLD time course in the precuneus and hippocampus. The present findings strongly suggest the critical role of the neurocognitive processes in these regions for situation model processing. They lend support to the idea that multi-sentence discourse processing draws on dynamic activations in a neural network that includes several regions beyond the classical language network (Ferstl et al. 2008; Binder 2016; Hagoort 2017; Hasson et al. 2018). The current evidence for the role of lateral parietal as well as medial parietal and frontal regions in shared situation model processing will be discussed in the following. In contrast, against our prediction, we did not find evidence for the right inferior frontal gyrus to play a strong role in situation model processing as implemented in the current design.

### 4.1. The role of medial prefrontal and parietal cortices in event schema processing and memory retrieval

In the present study, we found evidence for the ventromedial prefrontal cortex (vmPFC) to play an important role in situation model processing. Critically, in the vmPFC, the alignment of the BOLD time course between speakers and listeners was positively related to the listeners’ text comprehension, which is in line with previous findings with autobiographical narratives (Stephens et al. 2010). Moreover, the speaker-listener alignment was higher in the precuneus and hippocampus when the listeners subsequently showed good recall of text elements.

The vmPFC has previously been suggested to be a core region that allows for context-dependent processing of incoming information. It may participate in multiple memory-related cognitive processes that support gist-extraction and the inferencing of non-presented relationships (Gilboa and Marlatte 2017). In discourse processing more specifically, the vmPFC may play a role in activating an appropriate event schema and making it available to integrate currently incoming information into a coherent situation model of the discourse (see also, Nieuwenhuis and Takashima 2011; van Kesteren et al. 2012; Gilboa and Marlatte 2017; Robin and Moscovitch 2017). Interactions between the vmPFC and several medial temporal and basal structures (hippocampus, amygdala, ventral striatum) may be important in this process (Nieuwenhuis and Takashima 2011; van Kesteren et al. 2012; Milivojevic et al. 2015). Specific functions have been attributed to the different regions of this cortico-hippocampal network. For instance, Robin and Moscovitch (2017) propose that schematic representations are mediated by the vmPFC, gist-like representations by the anterior hippocampus and perceptually detailed, highly specific representations by the posterior hippocampus and neocortex. In a similar vein, Reagh and Ranganath (2018) argue that cortico-hippocampal connections underlie the reinstatement of specific event representations from memory. Accordingly, medial parietal and medial prefrontal regions are involved in constructing a situation model, informed by event schemas. The situation model is then populated with local features that are represented in anterior-temporal cortical areas. The hippocampus mediates and facilitates the integration of information between the two networks, which can sharpen their activity patterns into a representation of a specific event. Support for the idea of joint cortico-hippocampal roles in constructing situation models can also be found in research on event cognition. That is, over the course of a discourse, the situation model is updated on a local or on a global scale when new information is added. At event boundaries, the situation model is more likely to be updated globally. Brain regions that usually show sensitivity to such event transitions lie in the core of the default-mode network (e.g., medial parietal and medial frontal regions) as well as in the hippocampus, which are areas that also play a role in episodic memory retrieval (for a review, see Radvansky and Zacks 2017).

In the present study, the availability of a highly informative title provided a context in the form of an event schema. Even in the intermediately informative condition, the event schema was to a certain degree informative to allow for an integration of the incoming information. Yet a more controlled and effortful retrieval of world-knowledge information was required for building a conceptually coherent situation model. The present finding of increased BOLD responses and speaker-listener ISCs in medial frontal and parietal regions as well as the hippocampus when the title was informative lends support to the idea that not only situation model processing but also its sharing between comprehension and production critically draws on neural regions that are involved in schema activation and episodic memory retrieval. Importantly, the spatiotemporally similar neurocognitive activity between speakers and listeners underlying semantic convergence, schema activation, and episodic memory retrieval appears to constitute the key process allowing for the *parity of representations* between comprehension and production on the discourse level (Pickering and Garrod 2004; Pickering and Garrod 2006). The parity of representations relies on this shared part of the neurocognitive activity, even if the full conceptual processing may involve more widespread and differential activity in each of the interacting individuals.

### 4.2. The lateral parietal cortex as a high-level conceptual convergence zone

Beside the medial surface of the brain, the activation pattern observed for highly coherent discourse (highly informative title condition, HT) compared to the less coherent ones involved bilateral lateral parietal regions (angular gyrus, supramarginal gyrus). The present finding is in line with previous findings of a significant hemodynamic activation and ISC in these regions during coherent discourse-level processing (Martin-Loeches et al. 2008; Ames et al. 2015; Saalasti et al. 2019).

The lateral parietal cortex, together with inferior and middle temporal regions constitutes a network of high-level multimodal conceptual convergence zones (Binder and Desai 2011; Binder 2016) that is involved in constructing large-scale conceptual representations. The angular gyrus (AG), more specifically, has been suggested to function as a supramodal conceptual combination area (Menenti et al. 2008; Binder and Desai 2011; Humphreys and Lambon Ralph 2015; Price et al. 2015; Binder 2016; Hagoort 2017; Rugg and King 2018). This region plays a key role in remembering detailed experiences from the past but also in simulating vivid scenarios in the future (Ramanan et al. 2018). However, the AG is not a functionally homogenous region, but at least a dorsal (anterior) and a ventral (posterior) functional sub-region have been identified (Uddin et al. 2010). Interestingly, these AG sub-regions are characterized by different connectivity profiles, with the dorsal sub-region being more strongly connected with lateral prefrontal regions, the caudate nucleus and cingulate cortex, and the ventral sub-region more strongly with the precuneus, (para)hippocampal and medial frontal regions (Uddin et al. 2010; for a review, see also Ramanan et al. 2018). Moreover, in a recent meta-analysis, Humphreys and Lambon Ralph (2015) showed that the dorsal lateral parietal cortex plays a role in goal-directed, executive tasks, whereas the ventral lateral parietal cortex is consistently involved in automatic, stimulus-driven processes in verbal and non-verbal tasks. In the present study, situation model comprehension was associated with an involvement of both dorsal and ventral AG, but the involvement was strongest for the ventral part. Hence, establishing and updating a situation model may at least partially rely on the interaction of the ventral AG with (para)hippocampal regions, the precuneus and medial frontal regions. To sum up, neurocognitive processes of large-scale semantic and conceptual convergence in a network involving the lateral parietal cortex, notably the angular gyrus, seem to play an important role in the construction of a situation model and its sharing between speaker and listener.

### 4.3. The central role of neural hub regions in situation model processing

Strikingly, the regions beyond the classical language network that we found involved in the current task, namely the lateral parietal as well as medial prefrontal and parietal cortices and the hippocampus, overlap to a certain degree with parts of the semantic network (SN) and the theory of mind (ToM) network, but most importantly with parts of the default-mode (or task-negative) network (DMN; Fox and Raichle 2007; Raichle 2015). The DMN is a network that shows most hemodynamic activity during rest and is anti-correlated with activation in the dorsal attention network, i.e., the DMN deactivates during attention-demanding non-self-referential tasks (Fox and Raichle 2007; Humphreys et al. 2015). Previously, it has been argued that it may be inaccurate to consider the DMN as a mere *task-negative* network, not the least because of the numerous cognitive activities involved in resting and mind-wandering, which includes episodic memory retrieval, and semantic and social information processing (Binder and Desai 2011). This network seems to play an important role in various cognitive functions that are related to meaning processing, although with a functional differentiation between regions (Seghier and Price 2012). Findings of DMN regions showing strong activation in higher-order semantic and conceptual processing have led to the suggestion that some of the DMN regions, i.e., inferior parietal and inferior and middle temporal cortices, play an important role as high-level multi- or supramodal convergence zones (Binder and Desai 2011; Binder 2016). Moreover, this role in higher order conceptual processing could be based on these regions’ capacity to integrate information over longer timescales (Hasson et al. 2015; Simony et al. 2016). Importantly, this capacity may be linked to the fact that the DMN involves the most locally and globally connected hub regions of the brain. The connectivity profile of the DMN may partially underlie its function in multimodal integration and in processing spontaneous thoughts (Tomasi and Volkow 2011; van den Heuvel and Sporns 2013).

Furthermore, the DMN is not a functionally homogenous network but within the DMN, functional dissociations can be made. In a meta-analysis, Andrews-Hanna et al. (2014) identified a *medial temporal subsystem* (including the hippocampus, the parahippocampal cortex, the retrosplenial cortex (RSC), the posterior inferior parietal lobe, and the vmPFC) and a *dorso-medial subsystem* (including the dmPFC, the temporo-parietal junction, the lateral temporal cortex and the temporal pole), with the PCC and the anterior mPFC showing strong coherence with both subsystems. In this meta-analysis, the *medial temporal subsystem* was found to be functionally most involved in past and future autobiographical thought, episodic memory and contextual retrieval. In contrast, the *dorso-medial subsystem* was most involved in mentalizing and social cognition as well as story comprehension and semantic/conceptual processing. The core network that was shared between the two subsystems was activated during self-related processes, emotion/evaluation, and social and mnemonic processes (Andrews-Hanna et al. 2014). The present findings suggest that, due to their specific neurocognitive characteristics, parts of the DMN, especially the medial temporal subsystem, also play a role in situation model processing.

### 4.4. Conclusion

In the present study, discourse processing was associated with hemodynamic activation in the classical left-dominant fronto-temporal language network, but only the successful processing of a coherent discourse-level conceptual representation (situation model) additionally involved activation in the bilateral lateral parietal, medial parietal and prefrontal regions, as well as the hippocampus. The current data suggest that large-scale conceptual processing in discourse involves a neural network extending beyond classical language regions. Strikingly, this network of areas involved in meaningful discourse processing largely overlaps with hub regions in the default mode network. The coordination between regions involved in high-level conceptual convergence (e.g., lateral parietal cortex, middle temporal gyrus) and episodic memory retrieval (e.g., medial parietal, medial prefrontal, hippocampus) seems to be central to successful situation model processing in discourse. Dynamically changing situation model instantiations over the course of the discourse may thus be represented via a distributed neural network that changes with the incoming verbal information as well as with the information from memory, which needs to be integrated with discourse information. Critically, the spatiotemporally similar neurocognitive activity between speakers and listeners underlying semantic convergence and episodic memory retrieval appears to allow for the *parity of representations* between comprehension and production. In the present study, the evidence of ISC related to conceptual processing sheds further light onto the mechanisms underlying the capacity to share representations between interlocutors, which is fundamental for the communication of complex messages.

## 5. Acknowledgments

We are grateful to Marlou Rasenberg, Michel-Pierre Jansen, Maarten van den Heuvel, Birgit Knudsen and Iris Schmits for their help in the creation of the experimental material and in data coding, to Paul Gaalman and José Marques for their support in the setup of the fMRI experiment and to Roel Willems, Marloes Mak and Branka Milivojevic for fruitful discussions.

## References

Ames DL, Honey CJ, Chow MA, Todorov A, Hasson U. 2015. Contextual alignment of cognitive and neural dynamics. J Cogn Neurosci. 27:655–664.

Andersson JL, Jenkinson M, Smith S. 2007. Non-linear registration, aka Spatial normalisation FMRIB technical report TR07JA2. FMRIB Analysis Group of the University of Oxford. 2:1–21.

Andrews-Hanna JR, Smallwood J, Spreng RN. 2014. The default network and self-generated thought: component processes, dynamic control, and clinical relevance. Ann N Y Acad Sci. 1316:29–52.

Bakdash JZ, Marusich LR. 2017. Repeated Measures Correlation. Front Psychol. 8:456.

Baldassano C, Chen J, Zadbood A, Pillow JW, Hasson U, Norman KA. 2017. Discovering Event Structure in Continuous Narrative Perception and Memory. Neuron. 95:709–721 e705.

Bates D, Mächler M, Bolker B, Walker S. 2015. Fitting Linear Mixed-Effects Models Using lme4. J Stat Softw. 67.

Binder JR. 2016. In defense of abstract conceptual representations. Psychon Bull Rev. 23:1096–1108.

Binder JR, Desai RH. 2011. The neurobiology of semantic memory. Trends Cogn Sci. 15:527–536.

Boersma P, Weenink D. 2016. Praat: doing phonetics by computer. Version 6.0.16.

Boynton GM, Engel SA, Glover GH, Heeger DJ. 1996. Linear systems analysis of functional magnetic resonance imaging in human V1. J Neurosci. 16:4207–4221.

Bransford JD, Johnson MK. 1972. Contextual prerequisites for understanding: Some investigations of comprehension and recall. J Verbal Learning Verbal Behav. 11:717–726.

Brooks JC, Beckmann CF, Miller KL, Wise RG, Porro CA, Tracey I, Jenkinson M. 2008. Physiological noise modelling for spinal functional magnetic resonance imaging studies. Neuroimage. 39:680–692.

Cabeza R, Nyberg L. 2000. Imaging cognition II: An empirical review of 275 PET and fMRI studies. J Cogn Neurosci. 12:1–47.

Cavanna AE, Trimble MR. 2006. The precuneus: a review of its functional anatomy and behavioural correlates. Brain. 129:564–583.

Dodell-Feder D, Koster-Hale J, Bedny M, Saxe R. 2011. fMRI item analysis in a theory of mind task. Neuroimage. 55:705–712.

Dooling DJ, Lachman R. 1971. Effects of comprehension on retention of prose. J Exp Psychol. 88:216.

Ferstl EC, Neumann J, Bogler C, von Cramon DY. 2008. The extended language network: a meta-analysis of neuroimaging studies on text comprehension. Hum Brain Mapp. 29:581–593.

Fox MD, Raichle ME. 2007. Spontaneous fluctuations in brain activity observed with functional magnetic resonance imaging. Nat Rev Neurosci. 8:700–711.

Gilboa A, Marlatte H. 2017. Neurobiology of Schemas and Schema-Mediated Memory. Trends Cogn Sci. 21:618–631.

Gusnard DA, Akbudak E, Shulman GL, Raichle ME. 2001. Medial prefrontal cortex and self-referential mental activity: Relation to a default mode of brain function. Proc Natl Acad Sci U S A. 98:4259–4264.

Hagoort P. 2016. MUC (Memory, Unification, Control): A Model on the Neurobiology of Language Beyond Single Word Processing. In: Hickok G, Small SL, editors. Neurobiology of Language. San Diego: Academic Press p 339–347.

Hagoort P. 2017. The core and beyond in the language-ready brain. Neurosci Biobehav Rev. 81:194–204.

Hasson U, Chen J, Honey CJ. 2015. Hierarchical process memory: memory as an integral component of information processing. Trends Cogn Sci. 19:304–313.

Hasson U, Egidi G, Marelli M, Willems RM. 2018. Grounding the neurobiology of language in first principles: The necessity of non-language-centric explanations for language comprehension. Cognition. 180:135–157.

Hasson U, Nir Y, Levy I, Fuhrmann G, Malach R. 2004. Intersubject Synchronization of Cortical Activity During Natural Vision. Science. 303:1634–1640.

Hothorn T, Bretz F, Westfall P. 2008. Simultaneous inference in general parametric models. Biom J. 50:346–363.

Humphreys GF, Hoffman P, Visser M, Binney RJ, Lambon Ralph MA. 2015. Establishing task- and modality-dependent dissociations between the semantic and default mode networks. Proc Natl Acad Sci U S A. 112:7857–7862.

Humphreys GF, Lambon Ralph MA. 2015. Fusion and Fission of Cognitive Functions in the Human Parietal Cortex. Cereb Cortex. 25:3547–3560.

Jenkinson M, Bannister P, Brady M, Smith S. 2002. Improved optimization for the robust and accurate linear registration and motion correction of brain images. Neuroimage. 17:825–841.

Jenkinson M, Beckmann CF, Behrens TE, Woolrich MW, Smith SM. 2012. Fsl. Neuroimage. 62:782–790.

Jenkinson M, Smith S. 2001. A global optimisation method for robust affine registration of brain images. Med Image Anal. 5:143–156.

Kuperberg GR, Lakshmanan BM, Caplan DN, Holcomb PJ. 2006. Making sense of discourse: an fMRI study of causal inferencing across sentences. Neuroimage. 33:343–361.

Lam NH, Schoffelen J-M, Uddén J, Hultén A, Hagoort P. 2016. Neural activity during sentence processing as reflected in theta, alpha, beta, and gamma oscillations. Neuroimage. 142:43–54.

Logothetis NK, Pauls J, Augath M, Trinath T, Oeltermann A. 2001. Neurophysiological investigation of the basis of the fMRI signal. Nature. 412:150.

Martin-Loeches M, Casado P, Hernandez-Tamames JA, Alvarez-Linera J. 2008. Brain activation in discourse comprehension: a 3t fMRI study. Neuroimage. 41:614–622.

Menenti L, Petersson KM, Scheeringa R, Hagoort P. 2008. When Elephants Fly: Differential Sensitivity of Right and Left Inferior Frontal Gyri to Discourse and World Knowledge. J Cogn Neurosci. 21:2358–2368.

Milivojevic B, Vicente-Grabovetsky A, Doeller CF. 2015. Insight reconfigures hippocampal-prefrontal memories. Curr Biol. 25:821–830.

Nash JC, Varadhan R. 2011. Unifying Optimization Algorithms to Aid Software System Users: optimx for R. 2011. 43:14.

Nieuwenhuis IL, Takashima A. 2011. The role of the ventromedial prefrontal cortex in memory consolidation. Behav Brain Res. 218:325–334.

Northoff G, Bermpohl F. 2004. Cortical midline structures and the self. Trends Cogn Sci. 8:102–107.

Pickering MJ, Garrod S. 2004. Toward a mechanistic psychology of dialogue. Behav Brain Sci. 27:169–190, discussion 190-226.

Pickering MJ, Garrod S. 2006. Alignment as the Basis for Successful Communication. Res Lang Comput. 4:203–228.

Pickering MJ, Garrod S. 2013. An integrated theory of language production and comprehension. Behav Brain Sci. 36:329–347.

Price AR, Bonner MF, Peelle JE, Grossman M. 2015. Converging evidence for the neuroanatomic basis of combinatorial semantics in the angular gyrus. J Neurosci. 35:3276–3284.

R Core Team. 2014. R: A language and environment for statistical computing. Vienna, Austria: R Foundation for Statistical Computing.

Radvansky GA, Zacks JM. 2017. Event Boundaries in Memory and Cognition. Curr Opin Behav Sci. 17:133–140.

Raichle ME. 2015. The restless brain: how intrinsic activity organizes brain function. Philos Trans R Soc Lond B Biol Sci. 370.

Ramanan S, Piguet O, Irish M. 2018. Rethinking the Role of the Angular Gyrus in Remembering the Past and Imagining the Future: The Contextual Integration Model. Neuroscientist. 24:342–352.

Reagh ZM, Ranganath C. 2018. What does the functional organization of cortico-hippocampal networks tell us about the functional organization of memory? Neurosci Lett. 680:69–76.

Robin J, Moscovitch M. 2017. Details, gist and schema: hippocampal–neocortical interactions underlying recent and remote episodic and spatial memory. Curr Opin Behav Sci. 17:114–123.

Rugg MD, King DR. 2018. Ventral lateral parietal cortex and episodic memory retrieval. Cortex. 107:238–250.

Saalasti S, Alho J, Bar M, Glerean E, Honkela T, Kauppila M, Sams M, Jaaskelainen IP. 2019. Inferior parietal lobule and early visual areas support elicitation of individualized meanings during narrative listening. Brain Behav. 9:e01288.

Schoffelen J-M, Oostenveld R, Lam NHL, Uddén J, Hultén A, Hagoort P. 2019. A 204-subject multimodal neuroimaging dataset to study language processing. Scientific Data. 6:17.

Seghier ML, Price CJ. 2012. Functional Heterogeneity within the Default Network during Semantic Processing and Speech Production. Front Psychol. 3.

Sieborger FT, Ferstl EC, von Cramon DY. 2007. Making sense of nonsense: an fMRI study of task induced inference processes during discourse comprehension. Brain Res. 1166:77–91.

Simony E, Honey CJ, Chen J, Lositsky O, Yeshurun Y, Wiesel A, Hasson U. 2016. Dynamic reconfiguration of the default mode network during narrative comprehension. Nat Commun. 7:12141.

Smirnov D, Glerean E, Lahnakoski JM, Salmi J, Jaaskelainen IP, Sams M, Nummenmaa L. 2014. Fronto-parietal network supports context-dependent speech comprehension. Neuropsychologia. 63:293–303.

Smith SM. 2002. Fast robust automated brain extraction. Hum Brain Mapp. 17:143–155.

Stephens GJ, Silbert LJ, Hasson U. 2010. Speaker-listener neural coupling underlies successful communication. Proc Natl Acad Sci U S A. 107:14425–14430.

Tomasi D, Volkow ND. 2011. Association between functional connectivity hubs and brain networks. Cereb Cortex. 21:2003–2013.

Uddin LQ, Supekar K, Amin H, Rykhlevskaia E, Nguyen DA, Greicius MD, Menon V. 2010. Dissociable connectivity within human angular gyrus and intraparietal sulcus: evidence from functional and structural connectivity. Cereb Cortex. 20:2636–2646.

van den Heuvel MP, Sporns O. 2013. Network hubs in the human brain. Trends Cogn Sci. 17:683–696.

van Kesteren MT, Ruiter DJ, Fernandez G, Henson RN. 2012. How schema and novelty augment memory formation. Trends Neurosci. 35:211–219.

Wahlberg T, Magliano JP. 2004. The Ability of High Function Individuals With Autism to Comprehend Written Discourse. Discourse Process. 38:119–144.

Wiley J, Rayner K. 2000. Effects of titles on the processing of text and lexically ambiguous words: Evidence from eye movements. Mem Cogn. 28:1011–1021.

Winkler AM, Ridgway GR, Webster MA, Smith SM, Nichols TE. 2014. Permutation inference for the general linear model. Neuroimage. 92:381–397.

Wolfe MB, Magliano JP, Larsen B. 2005. Causal and semantic relatedness in discourse understanding and representation. Discourse Process. 39:165–187.

Xiang HD, Fonteijn HM, Norris DG, Hagoort P. 2010. Topographical Functional Connectivity Pattern in the Perisylvian Language Networks. Cereb Cortex. 20:549–560.

Zwaan RA, Radvansky GA. 1998. Situation models in language comprehension and memory. Psychol Bull. 123:162.

Zwaan RA, Singer M. 2003. Text comprehension. In: Graesser AC, Gernsbacher MA, Goldman SR, editors. Handbook of discourse processes. Hillsdale, NJ: Erlbaum p 83–121.

