## Supplementary Information for "No title, no theme: The joined neural space between speakers and listeners during production and comprehension of multi-sentence discourse"

### Materials and Methods

#### Pretest

In a pretest, an independent group of 30 Dutch native speakers (21 female, 9 male), between 18 and 32 years of age (average age: 23.1 ± 3.0 (SD) years) with no history of neurological, psychiatric or language disorders were asked to read and evaluate the 30 initial texts, from which 15 were subsequently chosen. Participants were presented with each text in one of the three conditions (highly informative title, intermediately informative title, no title) in a pseudorandomized order (max. three texts of one condition in immediate succession). Counterbalancing ensured that across participants, all texts were presented an equal number of times in each condition and that each participant was presented an equal number of texts per condition. Following the presentation of each title and text, the participants were asked to evaluate how easy it was to understand the text using a scale ranging from 1 ("I was totally confused") to 7 ("It was all totally clear"), then to identify the topic of the text and to indicate their confidence in their answer on a scale from 1 ("I am guessing randomly") to 7 ("I am totally certain"). Fifteen texts and their titles were considered appropriate for the subsequent fMRI experiment, if (1) the topic was not easily identifiable and no salient alternative topic emerged in the no title condition, and (2) there was a clear decreasing gradient of text comprehension scores for high (HT) > intermediate (IT) > no (NT) *Title informativeness*. Average text comprehension scores decreased with *Title informativeness* from high (HT): 5.49 ± 0.15 (SE), intermediate (IT): 4.25 ± 0.18 (SE), to none (NT): 3.42 ± 0.21 (SE) ($F(2,19)=32.2$, $p<.001$; Bonferroni-corrected pairwise comparisons: HT-IT: $p_{adj}<.001$; IT-NT: $p_{adj}<.001$; HT-NT: $p_{adj}<.001$; the statistical test applied in the pretest was identical to behavioral data analysis, for details see section 2.6.1).

Moreover, in order to obtain additional quality control for the intermediately informative title, the pretest involved a second part in which the participants were presented with the highly and intermediately informative title pairs for each text and were asked to evaluate (1) the degree of similarity of meaning and (2) the informativeness of the intermediately informative title. In the first part of the pretest, participants rated the degree of similarity of meaning of the two phrases on a scale ranging from 1 (no similarity of meaning) to 7 (perfect synonymy). For the 15 retained texts, the average assessed similarity was 4.52 ± 0.19 (SE). In the second part of the pretest, participants rated the informativeness of the intermediate title by indicating how easily they thought of the topic mentioned in the highly informative title when reading the intermediately informative title on a scale from 1 (not at all) to 7 (immediately). This measure is considered to indirectly address informativeness (or *semantic richness*), i.e., the number of shared (i.e., co-activated) semantic features (NoF; cf. Pexman et al. 2002) between the intermediately informative title and the highly informative title (i.e., the topic). For the 15 retained texts, the average assessed informativeness of the intermediately informative title was 3.78 ± 0.29 (SE) on a scale from 1 to 7.

**Localizer tasks**

After the main task, three functional localizer tasks were run in the following order: a *False belief localizer* (Dodell-Feder et al. 2011) in order to target the core theory of mind (ToM) network, a *WM/DMN (working memory/default mode network) localizer* and a *Language localizer* task (shortened version of Lam et al. 2016; Schoffelen et al. 2019). All localizer tasks were analyzed in the form of a block design. Six standard motion parameters were included in the statistical model. Functional regions-of-interest (fROIs) were identified using a group-level whole-brain *z*-map, family-wise error rate (FWE)-corrected at cluster-level $p<.05$ ($z$=3.1). A sphere of 10 voxels diameter was created around the peak coordinates. Selected fROIs are listed in the following after the description of each localizer task (see also Supplementary Table S 2 and Figure S 1).

1) *False belief localizer* (Dodell-Feder et al. 2011)

This task was run in order to target the core theory of mind (ToM) network (all bilateral: dmPFC, precuneus, inferior lateral parietal lobe), and more specifically top-down inferential rather than bottom-up perceptual ToM processes. Participants read 20 short stories, half of which were stories describing false beliefs and the other half were stories describing outdated (i.e., false) photographs and maps. Whereas in both conditions, participants were induced to represent false content, the conditions differed with respect to the type of false content represented (belief vs photograph/map). After each story, a true/false question that referred either to the situation in reality or to the false representation was presented. Participants responded to the question with a button response, i.e., using the right index finger to decide ‘true’ and the right middle finger to decide ‘false’. In the scanner, stories were presented visually for 10 s, followed by the true/false question for 4 s and finally 12 s of rest (a black screen). The duration of the *False belief localizer* task was of about 8 min. In order to localize the ToM network, the critical contrast was assessed as false beliefs > false photographs. Consistent with previous findings (Zaki and Ochsner 2012; Schurz et al. 2014), the selected ToM fROIs lay in the right medial prefrontal cortex [6/58/28], the left temporal pole [-56/4/-22], the (right) medial parietal cortex (precuneus [2/-56/36]) and in the right inferior parietal cortex (angular gyrus [58/-54/30]).

2) *WM/DMN (working memory/default mode network) localizer*

Specific contrasts in this localizer task were aimed at delineating fROIs of 1) verbal working memory, and 2) the default mode network (Fox and Raichle 2007; Raichle

2015). The *n*-back task is considered as a standard “executive” working memory (WM) task that requires participants to decide whether each stimulus in a sequence matches the one that appeared *n* items ago. The localizer task consisted of a blocked design alternating 2-back and 0-back variants of the n-back task using Dutch words and non-words. Words were bi-syllabic verbs taken from the texts that were presented in the main task (e.g., *werken* (to work), *weten* (to know), *drukken* (to press), *komen* (to come)). Non-words were verb-like pseudo-words that followed Dutch phonotactic constraints, created using Wuggy (Keuleers and Brysbaert 2010). Words and non-words were matched for number of syllables (2) and number of letters (6.0 ± 0.7 (SD)) and phonemes (4.5 ± 0.5 (SD)). Stimuli were presented in one run consisting of 30 blocks, i.e., 5 blocks per condition, presented in a pseudorandomized order. Within each block, all stimuli and the task were from the same condition (words/non-words; visual/auditory; 2-back/0-back) and blocks were presented in pseudorandomized order. In 2-back blocks, participants were asked to respond to every stimulus indicating whether or not the current word was the same as 2-back in the list (probability of the 2-back condition being satisfied (target) were 25% of the trials; 25% lure (n-1, n-3, n-4, n-5; 50% non-target non-lure)). In 0-back blocks, participants did a phoneme/letter-monitoring task, i.e., they were asked to respond to every stimulus indicating whether the current stimulus was a target (words/non-words that contain the target phoneme/letter; probability of stimulus being a target was 25% of the trials) or non-target (75% of words/non-words that do not contain target phoneme/letter). The target phoneme/letter (/b/, /k/, /l/, /s/, /v/) could occur in any position of the word. Participants were instructed to respond by pressing the right index finger button for ‘target’ items and the right middle finger button for ‘non-target’ items.

Each block started with an instruction about the modality of stimulus presentation (auditory/visual) and the task to perform (2-back/0-back target letter/phoneme), presented for 2 s. For phoneme monitoring, the target phoneme was auditorily presented in addition to the instruction screen figuring the orthographic representation of the target letter. For word stimuli, 2-back and 0-back blocks were presented in the visual and auditory modality whereas for non-word stimuli, only 0-back blocks (visual, auditory) were presented. Each block consisted of 12 stimuli. The duration of each block was 24 s, the interval between blocks 6 s (jittered between 4 and 8 s). The duration of the combined localizer task was approximately 16 min. Each trial consisted of 1000 ms presentation of a word or a non-word stimulus with an interstimulus interval of 1000 ms (jittered between 750 and 1250 ms), during which a fixation cross was presented in the center of the screen. The modality of presentation for half of the stimuli was visual (presented in the center of the screen), and the other half was auditory (fixation cross presented in the center of the screen). Within each block, the order of trials was pseudorandomly organized. Participants were instructed to respond as quickly and as accurately as possible. Feedback about response accuracy was given during the last 500 ms of the ITI in the form of a green (correct) or red (incorrect) cross in the place of the fixation cross in order to maximize the participants' attention. Before entering the scanner, participants did a practice run in order to familiarize with the task types.

The following contrasts were used to identify target fROIs for the main task: 1) verbal working memory (2-back words > 0-back words), and 2) the default mode network (0-back words > 2-back words). The selected verbal working memory fROIs lay in the right insula [36/20/-2], right superior parietal lobule (angular gyrus [38/-54/44]) and in the right middle/superior frontal gyrus [28/6/56], which reflected previous findings on especially the verbal component of working memory (Paulesu et al. 1993; Braver et al. 1997; Smith and Jonides 1999; Emch et al. 2019). The selected default mode fROIs lay in the left medial prefrontal cortex [-6/62/22], right hippocampus [26/-16/-16] and in the left medial parietal cortex (posterior cingulate [-4/-50/24]), which was in accordance with previous descriptions of the DMN (Fox and Raichle 2007; Andrews-Hanna et al. 2014; Raichle 2015).

3) *Language localizer* (shortened version of Lam et al. 2016; Schoffelen et al. 2019)

In this task, participants read individual words presented in the center of the screen, that in sequence constituted either (1) meaningful sentences or (2) random word lists. The sentences and word lists varied between 9 and 15 words in length. The word lists had been verified by a native speaker of Dutch so as not to allow for an emergence of coherent meaning. Six blocks per condition (*sentences, word lists*) were presented in alternation, each consisting of respectively 5 sentences or 5 word lists. For further detail about the stimulus material, see Lam et al. (2016). The duration of the language localizer task was about 12 min. The contrast sentences > word lists constituted the main contrast to localize the sentence processing network. In accordance with the literature (Xiang et al. 2010; Hagoort 2016, 2017), the selected sentence processing fROIs lay in the left middle/superior temporal gyrus [-56/-8/-12] and in the left inferior frontal gyrus [-54/20/22].

### Tables

**Table S 1. Titles for the 15 texts in the highly and intermediately informative title conditions.** The Dutch titles used in the experiment and their respective translations into English (italic font) are listed.

| Text | Highly informative title | Intermediately informative title |
|---|---|---|
| 1 | Een pompoen uitsnijden | Seizoen versiering voorbereiden |
| | *To carve a pumpkin* | *To prepare holiday decoration* |
| 2 | Op krukken lopen | Medische hulpmiddelen gebruiken |
| | *To walk on crutches* | *To use medical devices* |
| 3 | Een band vervangen | Een voertuig repareren |
| | *To change a tire* | *To repair a vehicle* |
| 4 | Kleren wassen | Een schoonmaak routine uitvoeren |
| | *To wash clothes* | *To carry out a cleaning routine* |
| 5 | Naaien | Precies handwerk uitvoeren |
| | *To sew* | *To do precision handicraft* |
| 6 | Muren schilderen | Een huis renoveren |
| | *To paint walls* | *To renovate a house* |
| 7 | Een sneeuwpop maken | Een figuur houwen |
| | *To build a snowman* | *To sculpt a figure* |
| 8 | Strijken | Stof verfraaien |
| | *To iron* | *To embellish tissue* |
| 9 | Paardrijden | Een sport beoefenen |
| | *Horseback riding* | *To do sports* |
| 10 | Diepte duiken | Watersport beoefenen |
| | *Deep diving* | *To do water sports* |
| 11 | Afwassen | Fragiele objecten schoonmaken |
| | *To wash dishes* | *To clean fragile objects* |
| 12 | Boogschieten | Precisie sporten beoefenen |
| | *Archery* | *To do precision sports* |

| | | |
|---|---|---|
| 13 | Bloemen water geven | Voor planten zorgen |
| | *To water flowers* | *To care for plants* |
| 14 | Koffie maken | Een drank bereiden |
| | *To make coffee* | *To prepare a beverage* |
| 15 | Naar de bioscoop gaan | Een vrije tijds activiteit doen |
| | *To go to the cinema* | *To do a leisure activity* |

**Table S 2. MNI coordinates for functional regions-of-interest (fROIs) obtained from the localizer tasks.** BOLD activation maxima from functional localizer contrasts that respectively target theory of mind (TOM), default mode (DMN), verbal working memory (WM) and language processing are listed.

| **Localizer** | **fROI** | **Label** | **Hemisph.** | **Region name** | **x** | **y** | **z** |
|---|---|---|---|---|---|---|---|
| ToM | TOM01 | rmPFC | R | medial frontal / prefrontal cortex | 6 | 58 | 28 |
| | TOM02 | lTP | L | temporal pole | -56 | 4 | -22 |
| | TOM03 | rPrec | R | medial parietal cortex / precuneus | 2 | -56 | 36 |
| | TOM04 | rAG | R | inferior parietal cortex / angular gyrus | 58 | -54 | 30 |
| WM | WM01 | rIns | R | insula | 36 | 20 | -2 |
| | WM02 | rSPL | R | superior parietal lobule / angular gyrus | 38 | -54 | 44 |
| | WM03 | rSFG | R | middle / superior frontal gyrus | 28 | 6 | 56 |
| DMN | DMN01 | lmPFC | L | medial frontal / prefrontal cortex | -6 | 62 | 22 |
| | DMN02 | rHPC | R | hippocampus | 26 | -16 | -16 |
| | DMN03 | lPCC | L | medial parietal cortex / posterior cingulate cortex | -4 | -50 | 24 |
| Language | LANG01 | lMTG | L | middle / superior temporal gyrus | -56 | -8 | -12 |
| | LANG02 | lIFG | L | inferior frontal gyrus | -54 | 20 | 22 |

Whole-brain *z*-maps were FWE-corrected at cluster-level $p<.05$ ($z=3.1$). Hemisph., Cerebral hemisphere.

### Figures

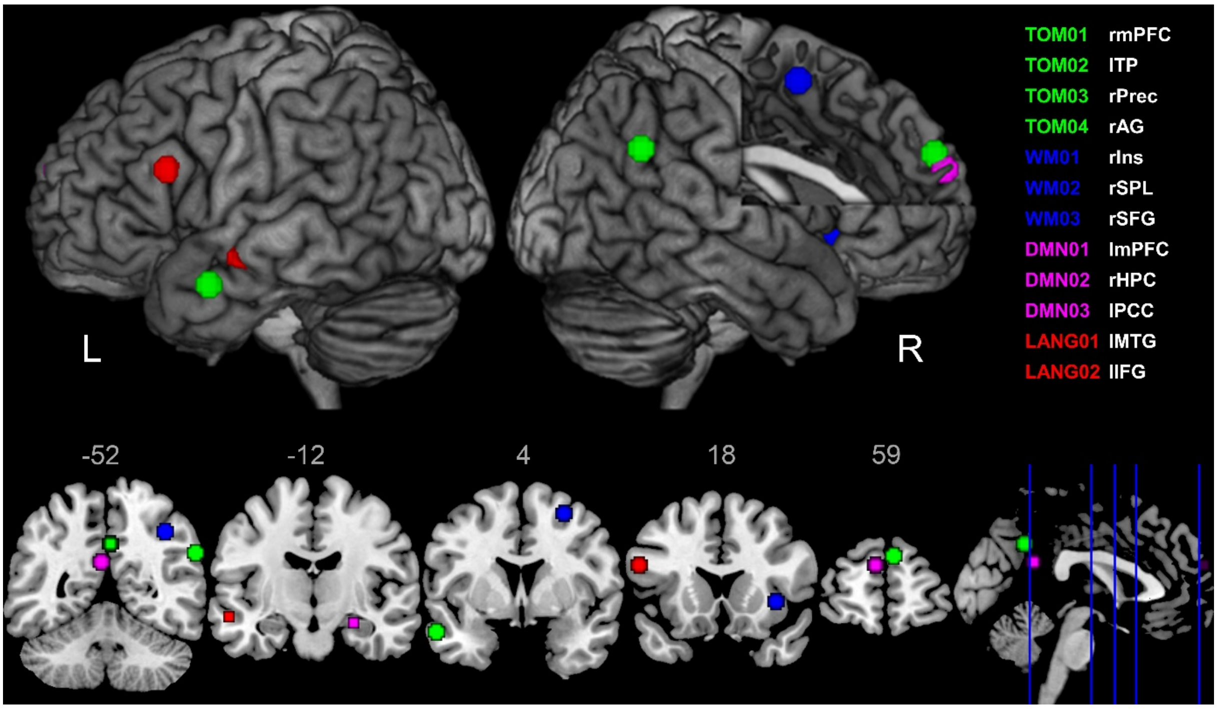


**Figure S 1. Functional regions-of-interest (fROIs) obtained from the localizer tasks.** BOLD activation maxima from functional localizer contrasts that respectively target theory of mind (TOM; green), default mode (DMN; violet), verbal working memory (WM; blue) and language (red) processing are shown. Spheres of 10 voxels diameter were created around the MNI coordinates of BOLD activation maxima. lIFG, left inferior frontal gyrus; lmPFC, left medial prefrontal cortex; lMTG, left middle temporal gyrus; lPCC, left posterior cingulate cortex; lTP, left temporal pole; rAG, right angular gyrus; rHPC, right hippocampus; rIns, right insula; rmPFC, right medial prefrontal cortex; rPrec, right precuneus; rSFG, right superior frontal gyrus; rSPL, right superior parietal lobule / angular gyrus.

**References**

Andrews-Hanna JR, Smallwood J, Spreng RN. 2014. The default network and self-generated thought: component processes, dynamic control, and clinical relevance. Ann N Y Acad Sci. 1316:29-52.

Braver TS, Cohen JD, Nystrom LE, Jonides J, Smith EE, Noll DC. 1997. A parametric study of prefrontal cortex involvement in human working memory. Neuroimage. 5:49-62.

Dodell-Feder D, Koster-Hale J, Bedny M, Saxe R. 2011. fMRI item analysis in a theory of mind task. Neuroimage. 55:705-712.

Emch M, von Bastian CC, Koch K. 2019. Neural correlates of verbal working memory: An fMRI meta-analysis. Front Hum Neurosci. 13:180.

Fox MD, Raichle ME. 2007. Spontaneous fluctuations in brain activity observed with functional magnetic resonance imaging. Nat Rev Neurosci. 8:700-711.

Hagoort P. 2016. MUC (Memory, Unification, Control): A Model on the Neurobiology of Language Beyond Single Word Processing. In: Hickok G, Small SL, editors. Neurobiology of Language. San Diego: Academic Press p 339-347.

Hagoort P. 2017. The core and beyond in the language-ready brain. Neurosci Biobehav Rev. 81:194-204.

Keuleers E, Brysbaert M. 2010. Wuggy: A multilingual pseudoword generator. Behav Res Methods. 42.

Lam NH, Schoffelen J-M, Uddén J, Hultén A, Hagoort P. 2016. Neural activity during sentence processing as reflected in theta, alpha, beta, and gamma oscillations. Neuroimage. 142:43-54.

Paulesu E, Frith CD, Frackowiak RS. 1993. The neural correlates of the verbal component of working memory. Nature. 362:342.

Pexman PM, Lupker SJ, Hino Y. 2002. The impact of feedback semantics in visual word recognition: Number-of-features effects in lexical decision and naming tasks. Psychon Bull Rev. 9:542-549.

Raichle ME. 2015. The restless brain: how intrinsic activity organizes brain function. Philos Trans R Soc Lond B Biol Sci. 370.

Schoffelen J-M, Oostenveld R, Lam NHL, Uddén J, Hultén A, Hagoort P. 2019. A 204-subject multimodal neuroimaging dataset to study language processing. Scientific Data. 6:17.

Schurz M, Radua J, Aichhorn M, Richlan F, Perner J. 2014. Fractionating theory of mind: a meta-analysis of functional brain imaging studies. Neurosci Biobehav Rev. 42:9-34.

Smith EE, Jonides J. 1999. Storage and executive processes in the frontal lobes. Science. 283:1657-1661.

Xiang HD, Fonteijn HM, Norris DG, Hagoort P. 2010. Topographical Functional Connectivity Pattern in the Perisylvian Language Networks. Cereb Cortex. 20:549-560.

Zaki J, Ochsner KN. 2012. The neuroscience of empathy: progress, pitfalls and promise. Nat Neurosci. 15:675-680.
